# Pairing Explainable Deep Learning Classification with Clustering to Uncover Effects of Schizophrenia Upon Whole Brain Functional Network Connectivity Dynamics

**DOI:** 10.1101/2023.03.01.530708

**Authors:** Charles A. Ellis, Robyn L. Miller, Vince D. Calhoun

## Abstract

Many studies have analyzed resting state functional magnetic resonance imaging (rs-fMRI) dynamic functional network connectivity (dFNC) data to elucidate the effects of neurological and neuropsychiatric disorders upon the interactions of brain regions over time. Existing studies often use either machine learning classification or clustering algorithms. Additionally, several studies have used clustering algorithms to extract features related to brain states trajectories that can be used to train interpretable classifiers. However, the combination of explainable dFNC classifiers followed by clustering algorithms is highly underutilized. In this study, we show how such an approach can be used to study the effects of schizophrenia (SZ) upon brain activity. Specifically, we train an explainable deep learning model to classify between individuals with SZ and healthy controls. We then cluster the resulting explanations, identifying discriminatory states of dFNC. We lastly apply several novel measures to quantify aspects of the classifier explanations and obtain additional insights into the effects of SZ upon brain network dynamics. Specifically, we uncover effects of schizophrenia upon subcortical, sensory, and cerebellar network interactions. We also find that individuals with SZ likely have reduced variability in overall brain activity and that the effects of SZ may be temporally localized. In addition to uncovering effects of SZ upon brain network dynamics, our approach could provide novel insights into a variety of neurological and neuropsychiatric disorders in future dFNC studies.

## INTRODUCTION

The analysis of resting state functional magnetic resonance imaging data (rs-fMRI) dynamic functional network connectivity (dFNC) has provided insight into a variety of neurological and neuropsychological disorders. Many studies involving dFNC have used machine learning classification models (1–9), and many have used clustering algorithms (10–28). A few studies have extracted features based on a clustering analysis and fed those into an interpretable machine learning model to identify key features related to disorders (6,11,18,29,30). However, the opposite approach of performing classification, applying explainability, and performing clustering presents a largely underutilized opportunity. Furthermore, it could easily be appended to many classification-based dFNC analyses for higher resolution insight into the effects of disorders. In this study, we build upon that opportunity to present a novel approach for the characterization of the effects of disorders upon brain network interactions. We apply our approach within the context of schizophrenia (SZ), first classifying between individuals with SZ and healthy controls (HCs), outputting explanations that identify the relative importance of dFNC features over time for the classification, and clustering the explanations to identify reoccurring states that have distinct differences in dFNC between individuals with SZ and HCs. We then extract several features to quantify how frequently SZs and HCs can be differentiated from one another based upon those features and how the effects of SZ upon dFNC vary over time and across brain networks. We uncover effects of SZ upon subcortical, sensory, and cerebellar network interactions. We also find that individuals with SZ likely have reduced variability in overall brain activity and that the effects of SZ may appear in a temporally localized manner. Our novel approach represents an advancement in the field of dFNC analysis and could be applied for enhanced insight into a variety of neurological and neuropsychological disorders in the future.

Several modalities are frequently applied for functional insight into neurological and neuropsychological disorders, including electroencephalography (EEG), magnetoencephalography (MEG), and fMRI. Moreover, all of these modalities have been used to study SZ (23,31–37). EEG and MEG offer high levels of temporal resolution. However, they have lower spatial resolution, which makes identifying the region of the brain from which signals originate challenging. In contrast, fMRI has a lower temporal resolution and a much higher spatial resolution. Importantly, fMRI represents a more common modality for SZ analysis than EEG, and particularly MEG. Both task (38,39) and resting-state (11,12,21,40–45) fMRI are used to study disorders. However, the majority of brain activity is spontaneous (i.e., reflects intrinsic networks that are not task modulated), which corresponds to the activity recorded in resting-state. Moreover, when individuals with a disorder perform a task, they may perform the task less effectively than healthy individuals, which introduces a potential confounder in any brain activity analyses (45).

In rs-fMRI analyses, a variety of features can be extracted to characterize different aspects of recordings. These include independent components (46) and spectral features (47)(1). Functional network connectivity (FNC) is a popular feature that provides insight into brain network interactions. There are two common forms of FNC - static FNC (sFNC) and dFNC – that have been applied for insight into a variety of disorders and cognitive functions, including major depressive disorder (2,48–50), SZ, Alzheimer’s disease (26,27,51), autism spectrum disorder (52), spatial orientation (28), and cognition (53). Early FNC analyses often used sFNC features, meaning that they analyzed the correlations between brain regions across whole recordings. Moreover, while some studies continue to use sFNC in a standalone manner (53,54) or in conjunction with dFNC (6,55,56), the use of dFNC to analyze how brain region interactions change over time has become much more common (6,13,23,24,29,41,48,52). This is largely due to how studies have shown that dFNC can provide greater insight into disorders than sFNC (57,58).

A number of studies have applied machine learning and deep learning-based classification approaches to dFNC, and some of those approaches have also combined explainability analyses with the classifiers to help characterize disorders by identifying the features used by classifiers to differentiate between individuals with disorders and healthy controls. These studies have sought to identify key effects of major depressive disorder (1)(2), fetal alcohol spectrum disorder (3), Alzheimer’s disease (4), and aging (5). One study sought to identify differences between SZ and bipolar disorder (6). Additionally, some studies have used other connectivity measures for insights into disorders like autism spectrum disorder (7). Lastly, while also providing insight into disorder dynamics, several dFNC classification studies have sought to provide insight into how deep learning classifiers might be used for clinical SZ diagnosis by developing approaches that combine explanations with estimates of confidence in explanations (8,9).

Although a number of studies have applied classification approaches to differentiate between individual with disorders and healthy controls, clustering represents another common analysis approach. These analyses often assign dFNC windows to states that can be used to summarize dFNC dynamics. Features can then be extracted from the state time-series and analyzed for further insight. Clustering analyses have been used to study the effects of early-stage SZ (10), the effects of SZ upon the default mode network (11,21,22), visual network (23), and whole-brain (24,25), the effects of Alzheimer’s disease upon the cognitive control network (26) and the whole-brain (27), and the relationship between whole-brain dFNC and spatial orientation (28). They have also been used to gain insight into differences in brain dynamics between individuals with SZ, schizoaffective disorder, and bipolar disorder (12). Due to the popularity of this analysis approach, numerous clustering approaches have been developed to further enhance the insights that can be obtained from these analyses (13–20).

Building upon the popularity of clustering approaches for dFNC analysis, a handful of studies have extracted features that summarize aspects of states identified during clustering and trained a classifier on those extracted features. These analyses have helped differentiate between healthy individuals, individuals with SZ, and individuals with bipolar disorder (6). They have been used to differentiate between SZs and healthy controls with default mode network dFNC (11,18) and whole-brain dFNC using the likelihood that individuals will transition between brain states (29). They have also been used in related modalities like EEG (30).

Given their extensive use in the field, both classification and clustering analyses have proved themselves to be highly useful and have provided many valuable insights. Nevertheless, they do have shortcomings. For example, in explainable classification, explanations are often averaged across a whole dataset or are examined on a global level. Given previous studies comparing local and global explanations (59), one can reasonably conclude that averaging or using global explanations could obscure the heterogeneous effects of disorders across individuals. Additionally, with the exception of a couple studies that quantify aspects of explanations (9,46), these studies often do not provide much insight into how disorders affect brain dynamics over time. Clustering analyses are very useful in providing insights into dynamics that would typically be obscured by classification approaches. However, as shown in studies that perform classification upon features extracted following clustering (6,18,29), the dynamical features identified by clustering are often not vary discriminatory (i.e., they yield low classification performance). The combination of an explainable classifier followed by a clustering approach presents an opportunity that could (1) improve upon clustering approaches by providing highly discriminatory insight into disorders and (2) improve upon classification approaches by providing more granular insights into the effects of disorders upon brain activity. Moreover, while we have applied a combination of a classifier with a clustering algorithm in one previous study to identify subtypes of SZs (60), the potential of this approach still remains largely understudied.

In this study, we present a novel approach that combines an explainable deep learning classifier with a subsequent clustering analysis that identifies discriminatory states of dFNC activity. We implement the approach to whole brain dFNC data from individuals with SZ and healthy controls. After identifying discriminatory states of dFNC activity, we then extract features from the identified state time-series to quantify how frequently SZs and HCs can be differentiated from one another based upon the dFNC patterns most important to each state. Lastly, we extract several features from the original classifier explanations to gain additional insight into the effects of SZ upon brain dynamics.

## METHODS

In this section, we describe our approach for the study. The approach is also shown in Figure 1. (1) We used a pre-existing dataset composed individuals with SZ (SZs) and healthy controls (HCs). (2) We preprocessed the data, extracting dFNC. (3) We trained a 1D-CNN to differentiate between HCs and SZs using whole-brain dFNC time-series. (4) We applied an explainability approach to identify key dFNC features differentiating HCs from SZs. (5) We clustered the explanations for each time step, identifying states in which HCs and SZs differed from one another, and (6) extracted features to quantify the amount of time that study participants spent in each state (i.e., occupancy rate, OCR) and the number of times that participants switched states (NST). (7) We next extracted features to quantify how the distribution of relevance between dFNC features varied over time. Lastly, we examined the relationship between the extracted features and (8) participant class and (9) SZ symptom severity. Note that the photo shown in step 1 of Figure 1 was taken from (61).

**Figure 1.**
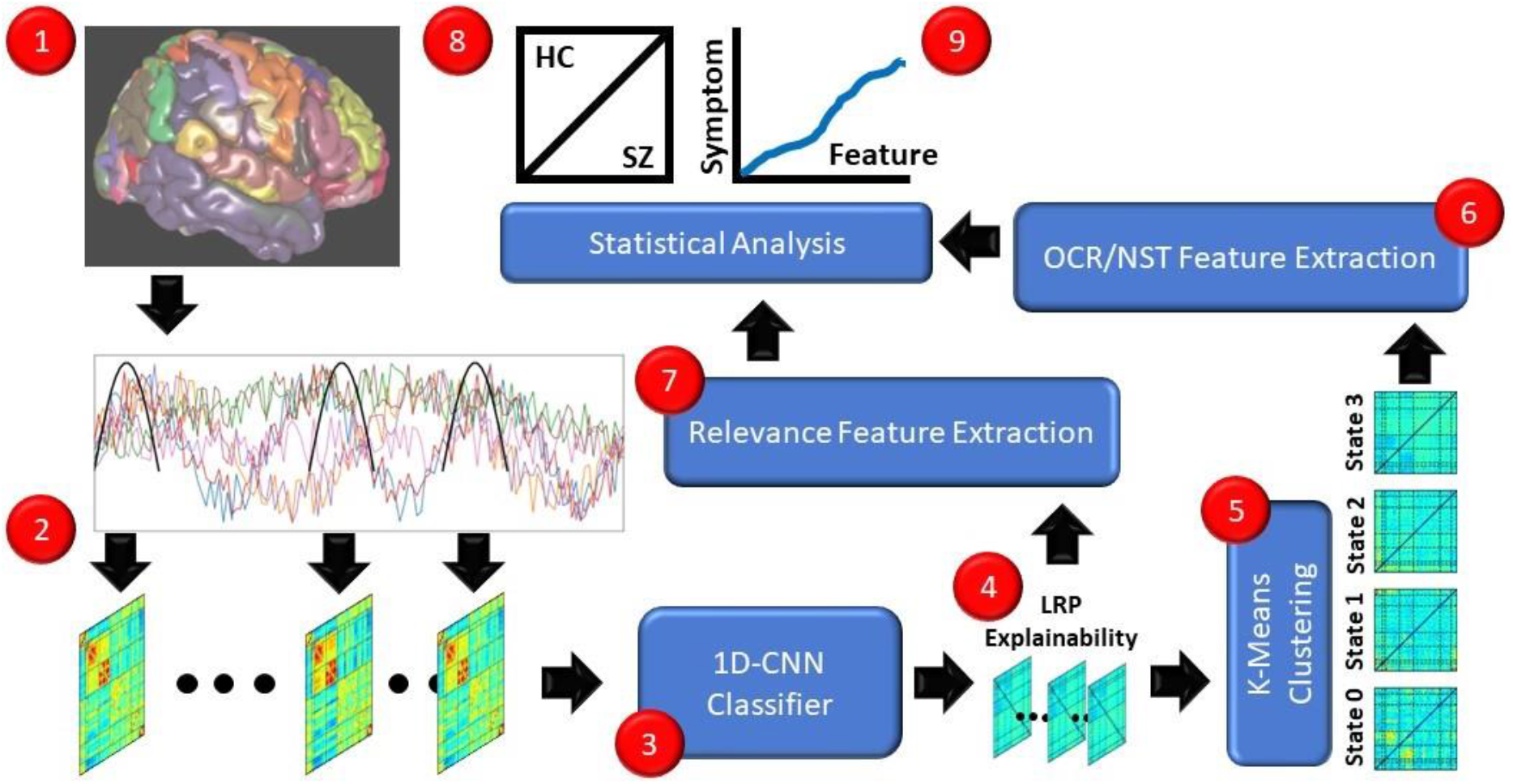
Methods Overview. (1) We used data composed of SZs and HCs. (2) We extracted whole brain dFNC. (3) We trained a 1D-CNN classifier using the dFNC time-series. (4) We applied Layer-wise Relevance Propagation to identify key features used by the classifier. (5) We applied k-means clustering to the explanations at each time step and identified several states. (6) We extracted features to quantify the amount of time that study participants spent in each state (OCR) and the number of times that participants switched states (NST). (7) We extracted relevance features to quantify how the distribution of relevance between dFNC features varied over time. Lastly, we examined the relationship between the extracted features and (8) class and (9) symptom severity.

### Description of Data

Table 1 shows the demographic and clinical information of study participants. We used rs-fMRI recordings from 160 HCs and 151 SZs that are part of the Functional Imaging Biomedical Informatics Research Network (FBIRN) dataset (62). The dataset has been used in many studies (8,9,11,18,22,54,60,63) and also contains negative and positive symptom scores from the Positive and Negative Syndrome Scale (PANSS) (64). Negative SZ symptoms include apathy, alogia, asociality, and affective flattening, while positive SZ symptoms include delusions, hallucinations, and bizarre behavior (41). The dataset was collected at 7 sites, including: Duke University/the University of North Carolina at Chapel Hill, the University of Iowa, the University of Minnesota, the University of California at San Francisco, the University of California at Los Angeles, the University of California at Irvine, and the University of New Mexico. All study participants provided written informed consent, and all data collection procedures were approved by the institutional review boards at each study site. Six study sites used 3T Siemens TIM Trio Scanners, and the remaining site used a 3T GE MR750 scanner. A T2*-weighted AC-PC aligned echo-planar imaging (EPI) sequence was used during data collection (TR = 2s, TE = 30ms, voxel size = 3.4 x 3.4 x 3.4 mm^3^, flip angle = 77°, slice gap = 1mm, number of frames = 162, and acquisition time = 5 min and 38s).

**Table 1.**
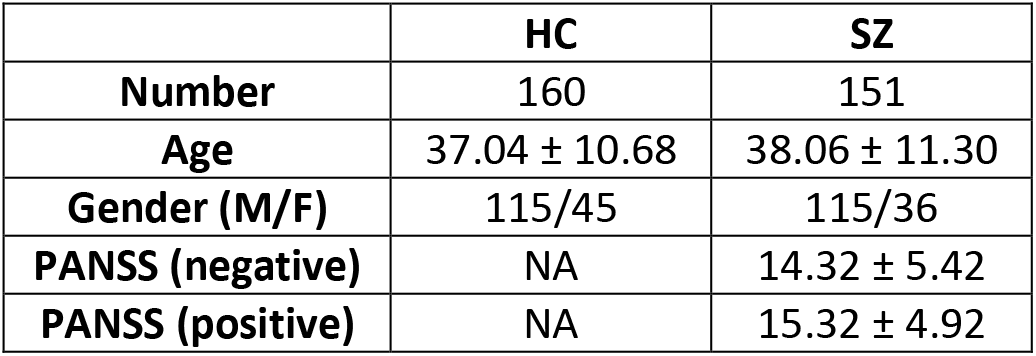
Participant Demographic and Clinical Information

|  | **HC** | **SZ** |
| --- | --- | --- |
| **Number** | 160 | 151 |
| **Age** | 37.04 ± 10.68 | 38.06 ± 11.30 |
| **Gender (M/F)** | 115/45 | 115/36 |
| **PANSS (negative)** | NA | 14.32 ± 5.42 |
| **PANSS (positive)** | NA | 15.32 ± 4.92 |

### Description of Data Preprocessing

The first five mock scans were removed prior to preprocessing, and preprocessing was conducted with statistical parametric mapping (SPM12, https://www.fil.ion.ucl.ac.uk/spm/). Rigid body motion correction was used to correct for head motion, and the data was spatially normalized to an EPI sequence template in the Montreal Neurological Institute (MNI) space. Images were then resampled to 3×3×3 mm^3^ and were smoothed via the application of a Gaussian kernel with a 6mm full width at half maximum. We then applied the Neuromark Pipeline of the Group Independent Component Analysis (ICA) of fMRI Toolbox (GIFT, http://trendscenter.org/software/gift). Neuromark uses spatially constrained ICA to extract components (ICs) reproducibly across datasets while also allowing the components to adapt to each dataset individually. We extracted 53 components that had peak activations in the gray matter of 7 brain networks using the neuromark_fMRI_1.0 template. The 53 components can be assigned to the subcortical (SCN, 5 ICs), sensorimotor (SMN, 9 ICs), auditory (ADN, 2 ICs), visual (VSN, 9 ICs), default mode (DMN, 7 ICs), cognitive control (CCN, 17 ICs), and cerebellar (CBN, 4 ICs) networks. We next extracted dFNC by calculating Pearson’s correlation between each pair of ICs using a sliding tapered window. The tapered window was formed by convolving a rectangle with a 40-second step size with a Gaussian with a standard deviation of 3. The choice of step size does affect findings. However, a 40-second step size has been used in a variety of studies (11,22,23,65). The resulting data was composed of 1,378 dFNC features and 124 time points per participant. Throughout the remainder of the study dFNC features corresponding to the interaction of particular networks will be abbreviated as Network1/Network2 (e.g., DMN/SMN). It should also be noted that interactions between any given pair of networks are identical if their order is switched (i.e., DMN/SMN = SMN/DMN).

### Description of Model Development

Figure 2 shows our 1D-CNN model architecture, which we originally used in (60). Before training, we feature-wise z-scored the dFNC data for each participant separately. The input data was of dimensions 124 time points x 1,378 features, The model was trained using Keras 2.2.4 (66). We trained the model using a 10-fold stratified shuffle split cross-validation approach. Training, validation, and test sets were composed of 80%, 10%, and 10% of the data, respectively. We used data augmentation to increase the size of the training set by a factor of 4. The augmentation involved creating 3 copies of the training data and adding Gaussian noise with a mean of zero and standard deviations of 0.7, 0.5, and 0.2 depending upon the copy. To alleviate the potential effects of class imbalances, we used a class-weighted categorical cross-entropy loss function. We used the Adam optimizer (67) with an adaptive learning rate starting at 0.001 and decreasing by 50% if 15 epochs passed without an increase in validation accuracy (ACC). We used a batch size of 50 and trained the model for 75 epochs. The first layer used Glorot uniform initialization (68), and all other layers used Kaiming He normal initialization (69). We performed testing using the models that corresponded to the training epochs with the highest validation accuracy, and we evaluated model test performance by calculating the mean and standard deviation of the ACC, sensitivity (SENS), and specificity (SPEC) across folds.

**Figure 2.**
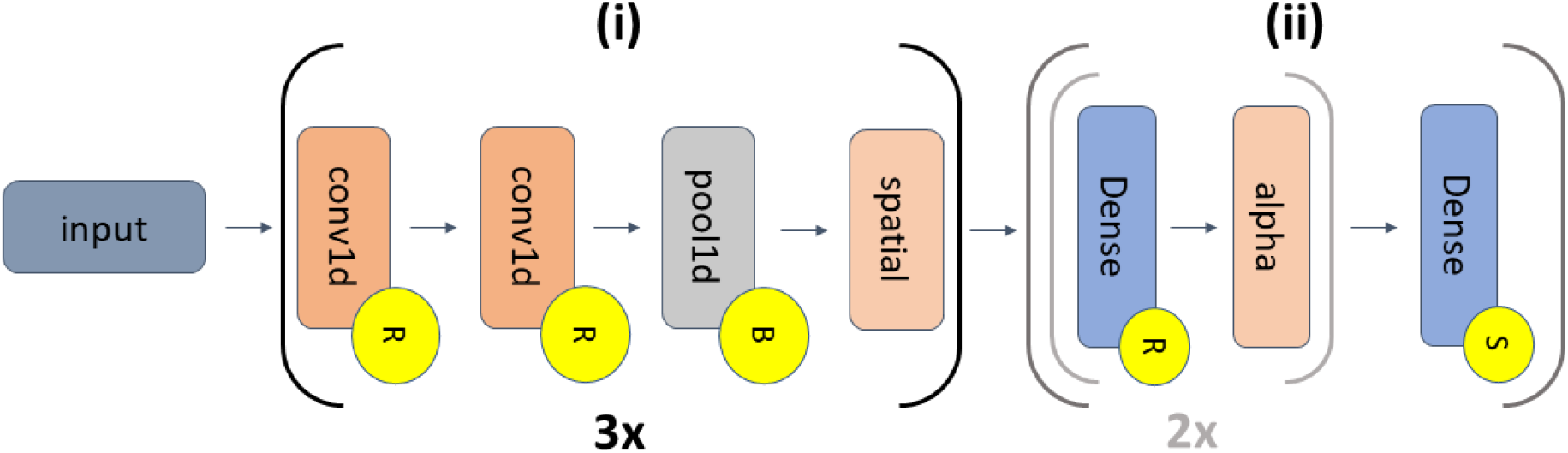
1D-CNN Architecture. The model has 2 components: (i) feature extraction (which repeats 3 times) and (ii) classification. The grey selection of (ii) repeats twice. The 3 pairs of convolutional (conv1d) layer pairs have 16, 24, and 32 filters, respectively (kernel size = 4). The pairs are followed by max pooling (pool size = 2) and spatial dropout (spatial, rates = 0.15, 0.25, and 0.3). Component (ii) has 3 dense layers (18, 14, and 2 nodes). The first two dense layers have alpha dropout (alpha) with rates of 0.35 and 0.25. ReLU activations, batch normalization, and softmax activations are represented by yellow circles containing an “R”, “B”, or “S”.

### Description of Explainability Approach

For explainability, we used the αβ-rule (70) of layer-wise relevance propagation (LRP) (71,72). LRP is a popular approach that have been used in many studies for insight into neurological time-series and neuroimaging data (8,9,59,60,73–80). LRP involves several steps. (1) A sample is forward passed through a network. (2) A total relevance of 1 is assigned to the output node of interest. (3) The total relevance is propagated back through the network using a relevance rule such that the total relevance is conserved from layer to layer until relevance is propagated to the input data space. Depending upon the relevance rule, both positive and negative relevance can be propagated through the network. Positive relevance highlights features that support the assignment of a sample to the class of interest, and negative relevance highlights features that support the assignment of the sample to the opposite class. To filter negative relevance and only propagate positive relevance, we used the αβ-rule with (α = 1, β = 0), which is shown in the equation below.

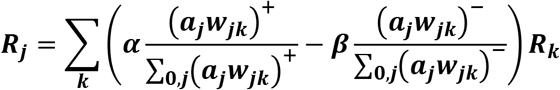

Where the subscripts *j* and *k* correspond to values for one of *J* nodes in a shallower layer and one of *K* nodes in a deeper layer, respectively. The shallower layer activation output is *a_j_*, and w refers to the model weights. The variables α and β affect positive and negative relevance propagation, respectively.

We used LRP to extract relevance for each training sample across folds. After extracting relevance, we calculated the absolute relevance of each sample and normalized them such that the total absolute relevance for each sample summed to exactly 1. While LRP is theoretically supposed to maintain a total relevance of 1 across all layers, practically that is not always the case. Our normalization of the relevance enabled an easier comparison across samples.

### Description of Clustering Approach

After outputting the relevance for each participant, we averaged the relevance of each participant across folds. We next normalized each time step such that the relevance at each time step summed to one and concatenated the relevance output for all training participants. After processing the relevance values, we applied k-means clustering with Euclidean distance and 500 initializations to assign the relevance value for each time step to a cluster. We swept from 2 to 10 clusters and selected the optimal number of clusters based on the silhouette score (81). We visualized the centroids for each cluster (i.e., state) and visualized the mean dFNC for time steps assigned to each step.

### Description of Clustering Feature Extraction

After assigning each relevance time step to a state, we created an array of state trajectories, such that each participant transitioned across states over time. Using the state trajectories, we calculated two sets of dynamical features to summarize aspects of the time-series. (1) We calculated the percent of time that each participant spent in each state (i.e., the occupancy rate, OCR), and (2) we calculated the number of state transitions (i.e., the number of times participants switched states, NST). Although, this is the first instance of applying these metrics to relevance time-series, both the OCR and NST have been used in a number of studies focused on clustering dFNC time-series (23,26–28).

### Description of Relevance Feature Extraction

We next analyzed the original participant-level normalized relevance independent of clustering to examine different aspects of the distribution of the relevance. (1) We summed the relevance across dFNC features for each participant. We then normalized the relevance of each participant over time (i.e., so that it summed to 1) and analyzed the temporal concentration of relevance by calculating the Earthmover’s or Wasserstein distance (EMD) between the normalized relevance and a uniform distribution of relevance (i.e., total relevance divided by number of time steps). EMD has been used in several fMRI dFNC studies (8,9,18,46). (2) We examined how greatly the distribution of relevance across all dFNC features shifted over time. To this end, we normalized the relevance at each time point such that the relevance across features at each time point summed to 1. We then calculated the Kullback-Leibler divergence (KLD) between each consecutive time step. We calculated the mean, median, range, and standard deviation of the KLD across time steps for each participant. One other study has used KLD for insight into dFNC (18). However, to our knowledge, no other studies have applied KLD to summarize relevance distributions.

### Description of Class-Level Statistical Analysis

We wanted to determine whether there were significant class-level differences in the OCR, NST, EMD, and KLD features. As such, we performed a series of two-tailed t-tests comparing the features for SZs and HCs. We analyzed the resulting t-statistics for insight into the directionality of differences between HC and SZ features. Additionally, to reduce the likelihood of false positives, we applied the Benjamini-Hochberg method for false discovery rate (FDR) correction (82) separately to p-values of the OCR and KLD features.

### Description of Symptom Severity Analysis

Lastly, we sought to understand whether there were relationships between the OCR, NST, EMD, and KLD features and the positive and negative PANSS scores. To this end, we performed ordinary least squares regression with age, gender, and negative PANSS or age, gender, and positive PANSS as independent variables and the features as dependent variables. This enabled us to control for the effects of age and gender. We then applied FDR correction (82) to the OCR and KLD p-values for each symptom score.

## RESULTS

In this section, we describe the performance results of our classifier, our classifier explanations before and after clustering, class-level differences in the features we extracted to quantify state trajectory dynamics, and class-level differences in the features that we extracted to quantify aspects of the relevance time-series.

### Differentiating Between HCs and SZs with a Deep Learning Classifier

Given the application of the explainability approach to the training data, it is important to note that model training performance tended to be near 100%. Additionally, all classifier test performance metrics were well above chance-level. Model SENS, SPEC, and ACC were 76.88 ± 10.84, 81.88 ± 7.1, and 79.38 ± 5.45, respectively. The model SPEC had a higher mean and lower standard deviation than the model SENS. Relative to dFNC classifiers using the FBIRN dataset in other studies, our model performed slightly better across all metrics (9,63).

### Identifying States of Differentiation between HCs and SZs

Using the silhouette method, we determined that 2 clusters, followed by 4 clusters, was the optimal number of clusters. As such, out of a desire to ensure that clusters were not simply an assignment of each class to separate clusters, we used 4 clusters in our analysis. Figure 3 shows the median dFNC and median normalized relevance per class. Figures 4, 5, and 6 show the relevance centroids for each state, the median relevance per class and state, and the median dFNC per class and state, respectively. Additionally, Table 2 shows the median percent of unnormalized relevance for samples in each class assigned to each state to give insight into the relative importance of samples in each state to the overall classification.

**Figure 3.**
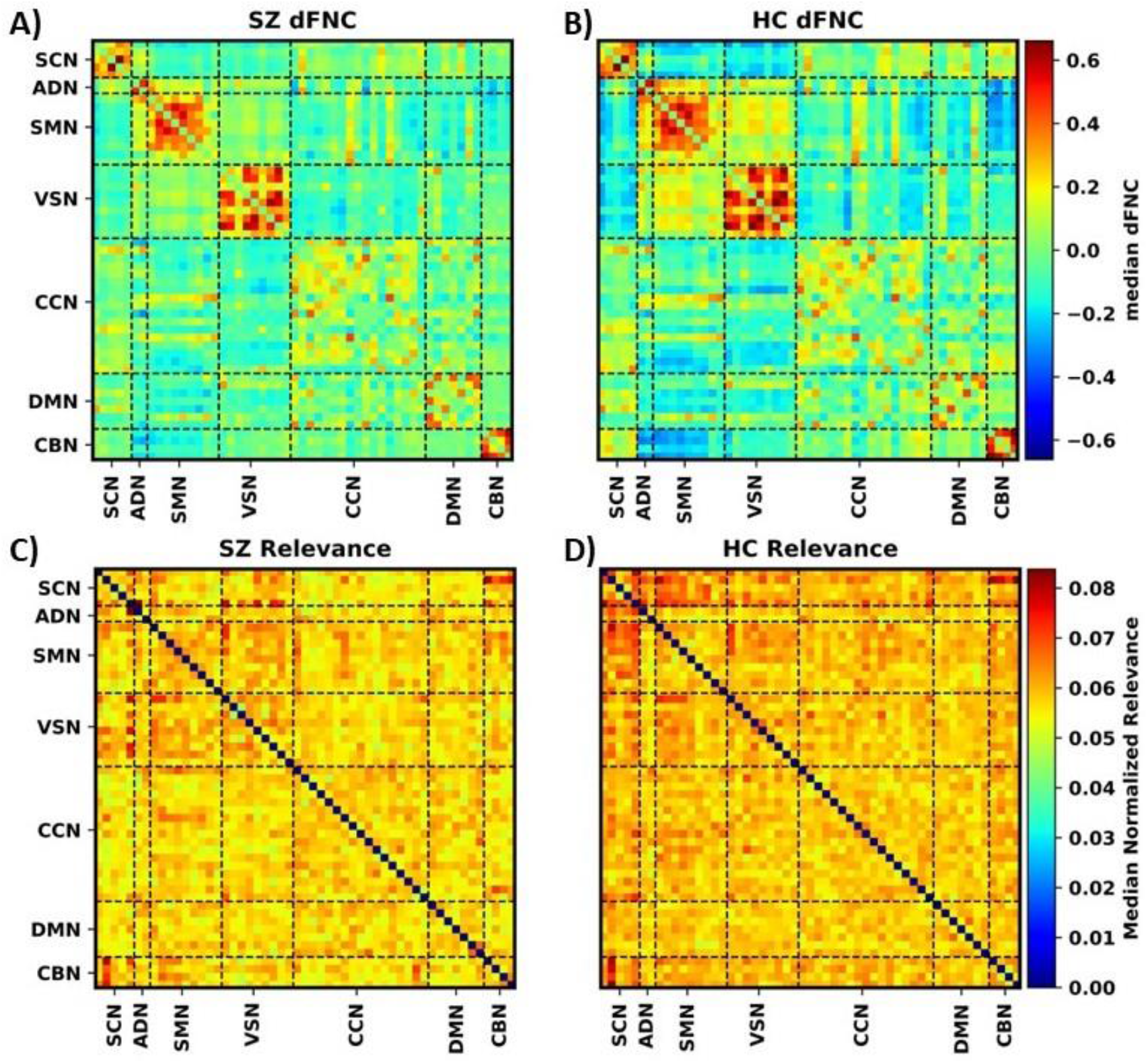
Mean dFNC and Relevance Per Class. Panels A) and B) show the mean dFNC for SZs and HCs, respectively, and share the same color bar to the right of the panels. Panels C) and D) show the mean relevance for SZs and HCs, respectively, and share the same color bar to the right of the panels.

**Figure 4.**
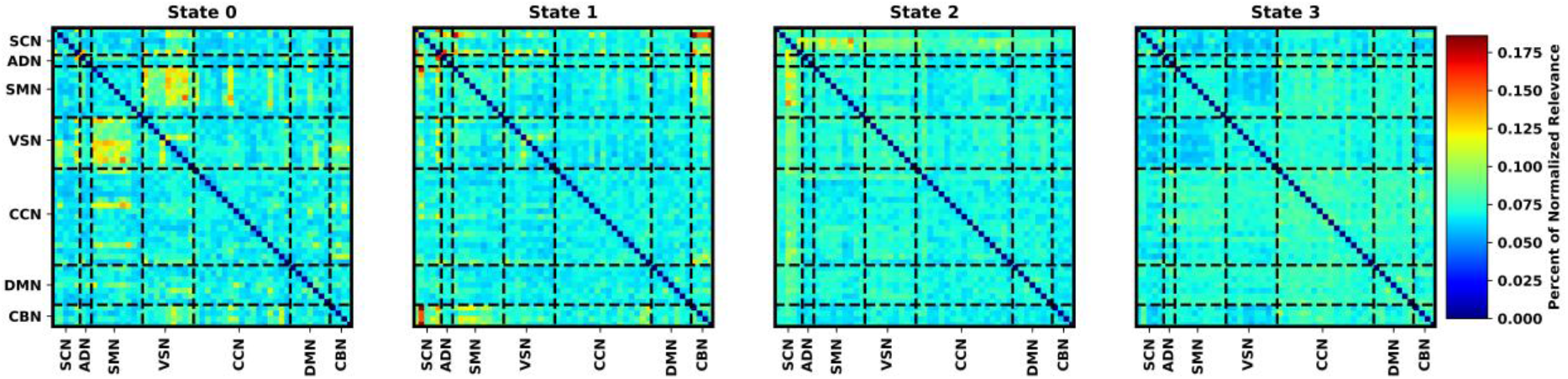
Relevance Cluster Centroids. Each panel shows the centroid for a different cluster (i.e., state) identified via k-means. The state that corresponds to each panel is shown in the panel title. All panels share the same color bar to the right of the figure. The centroids are displayed as correlation matrices, where each network is shown on the x- and y-axes. Black dashed lines separate each domain pair.

**Figure 5.**
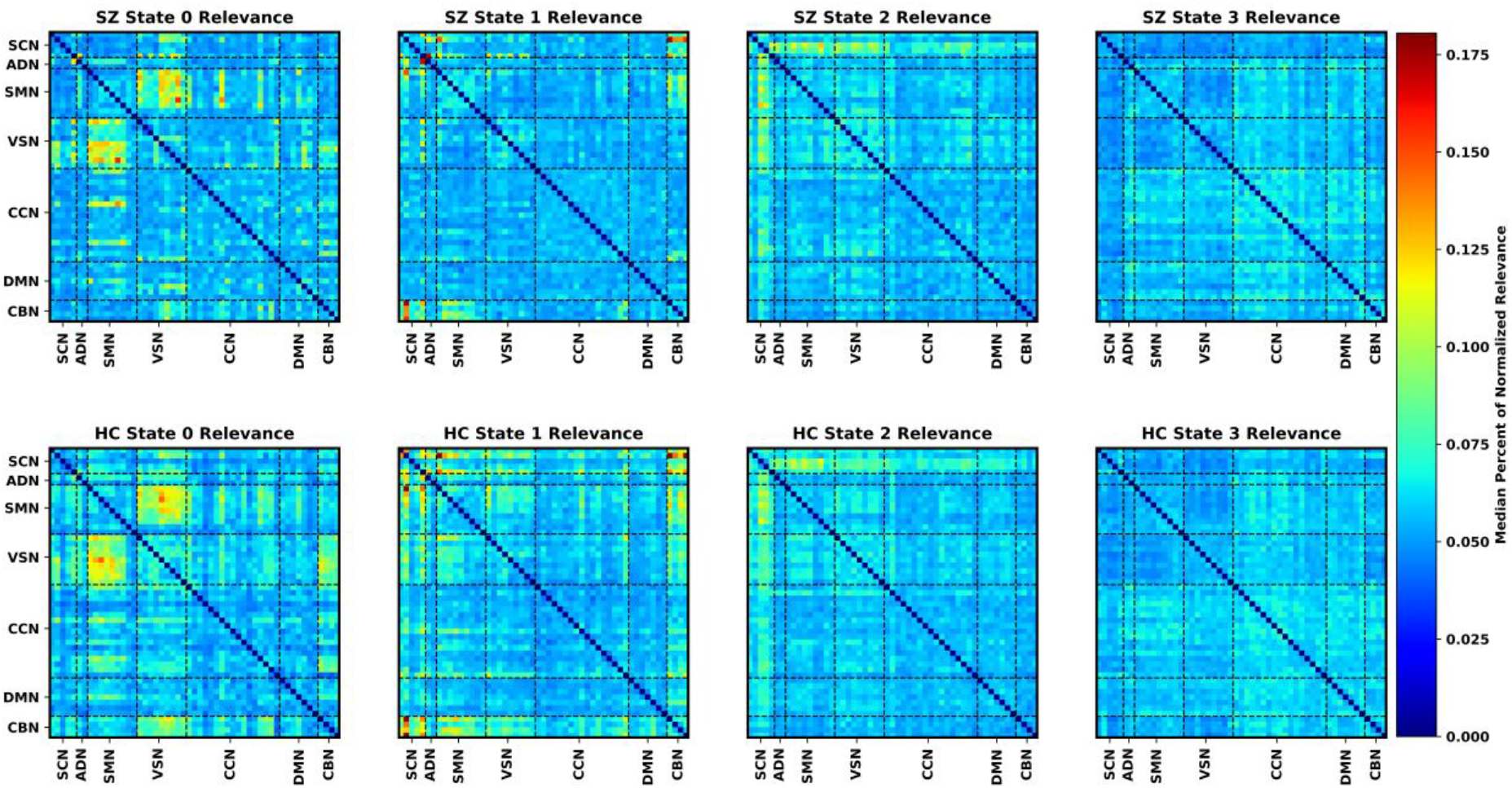
Median Relevance Per State and Class. Each panel shows the median relevance for each identified state and class. Note that the relevance for each time step was not normalized in this figure. SZs and HCs are shown in the top and bottom row of panels, respectively. States 0 through 3 are in ascending order from left to right. All panels share the same color bar to the right of the figure. The centroids are displayed as correlation matrices, where each network is shown on the x- and y-axes. Black dashed lines separate each domain pair.

**Figure 6.**
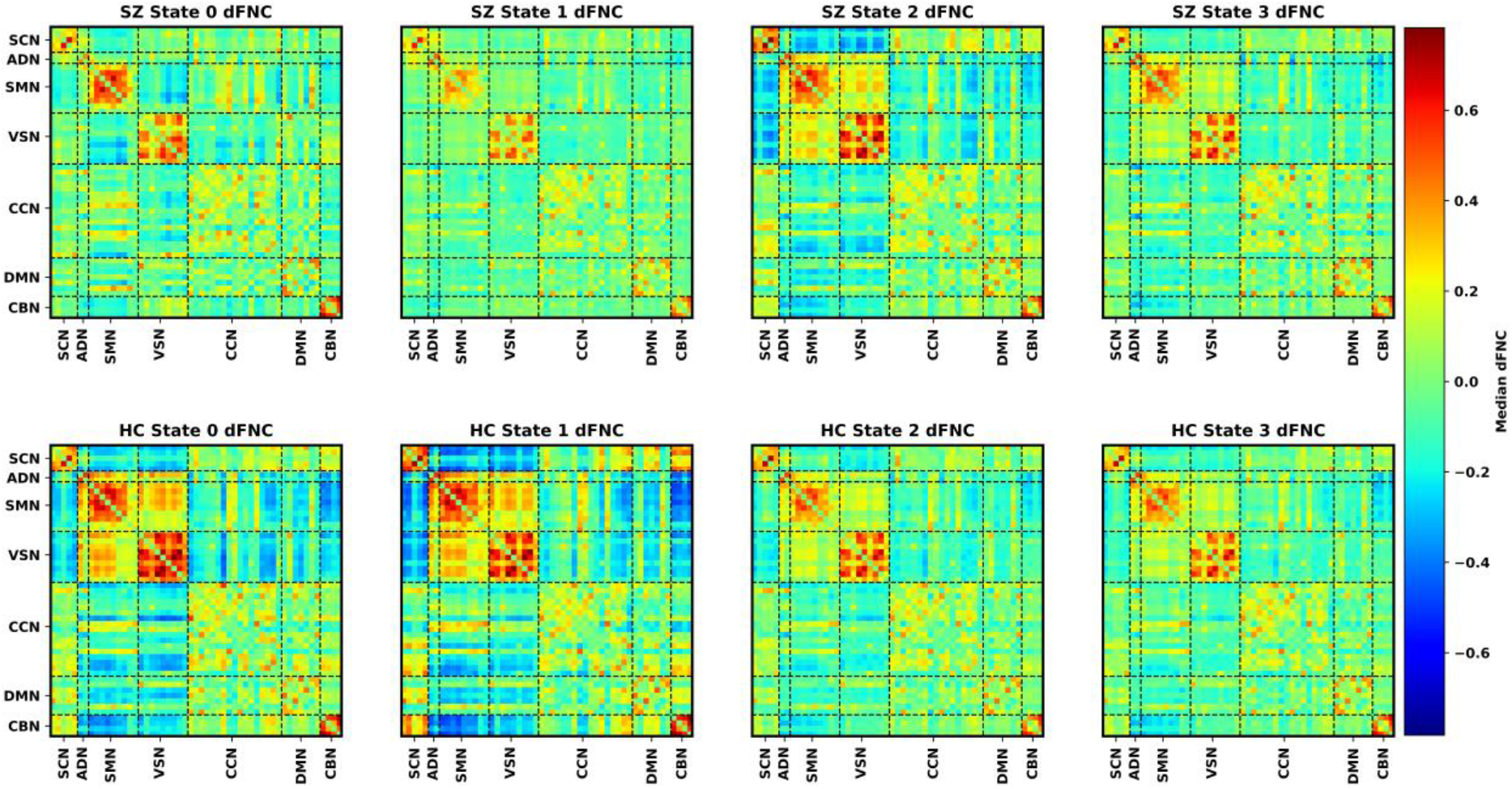
Median dFNC Per State and Class. Each panel shows the median dFNC for each identified state and class. SZs and HCs are shown in the top and bottom row of panels, respectively. States 0 through 3 are in ascending order from left to right. All panels share the same color bar to the right of the figure. The centroids are displayed as correlation matrices, where each network is shown on the x- and y-axes. Black dashed lines separate each domain pair.

**Table 2.** Median Unnormalized Relevance Per State and Class

|  | **State 0** | **State 1** | **State 2** | **State 3** |
| --- | --- | --- | --- | --- |
| **SZ** | 0.802 | 0.798 | 0.545 | 0.625 |
| **HC** | 0.664 | 0.867 | 0.701 | 0.676 |

Based on Figure 3, it seems that HCs have slightly more spatially concentrated relevance than SZs (i.e., the median relevance values are larger). Additionally, it seems that there are differences in importance in intra-SCN, SCN/VSN, and SCN/SMN connectivity between HCs and SZs. As shown in Figures 4 and 5, state 0 tends to prioritize VSN/SCN, VSN/SMN, and VSN CBN interactions. Additionally, parts of the VSN are more important for HCs than SZs. Relative to other states, state 1 tends to prioritize CBN/SCN, CBN/SMN, and CBN/VSN interactions. VSN interactions with part of the SMN are also important. HC classification also tends to place more importance upon CBN/VSN state 1 interactions than SZ classification. State 2 prioritizes intra-SCN interactions and interactions between the SCN and all other networks more than all other states. It prioritizes different parts of the VSN for SCN/VSN interactions than states 0 and 1. Additionally, median relevance distributions for state 2 HCs and SZs are very similar. State 3 relevance tends to be slightly more diffuse. However, it tends to focus more on CCN and DMN interactions with all other networks. As shown in Table 2, median unnormalized relevance tends to be slightly lower in SZs than HCs in all states except state 0, indicating that relevance for SZs tends to be a bit more diffuse. State 1 was most important for identifying HCs by a wide margin, followed by state 2, state 3, and state 0. State 0 was most important for identifying SZs, followed narrowly by state 1. State 2 and 3 were also important but much less so.

Based on Figure 3, there are visible differences in SCN/VSN, SCN/SMN, and CBN/SMN dFNC. There also seem to be slight differences in parts of CCN/SMN and CCN/VSN connectivity. As shown in Figure 5, state 0 SZ dFNC tended to be differentiated from the median SZ and HC dFNC of other states by its negative VSN/SMN activity, and state 0 HC dFNC tended to be differentiated from the median dFNC of other states by having moderately negative VSN/SCN and VSN/CBN activity. State 1 SZ dFNC is differentiated from SZ dFNC and HC dFNC in other states by having generally low magnitude connectivity. State 1 HC dFNC is more strongly differentiated from other HC states by having strongly negative CBN/SMN and CBN/VSN connectivity and moderately positive CBN/SCN connectivity. Additionally, median state 1 HC dFNC was very different from all SZ state median dFNC values. While it is similar in the directionality of the dFNC to SZ state 2, it has greater dFNC magnitudes. State 2 SZ dFNC is differentiated from SZ dFNC in other states by having more strongly positive intra-SCN and negative SCN/SMN and SCN/VSN activity. It is more like the HC dFNC medians than the other SZ states. State 2 HC dFNC is different from other HC dFNC states by having slightly less strongly negative SCN/VSN connectivity. State 3 HC and SZ dFNC tends to be similar to other states. However, CCN/SMN, CCN/VSN, DMN/SMN, and DMN/VSN connectivity tends to be slightly positive in magnitude in contrast to states 0 and 1 HC dFNC.

### Quantifying Class-Level Differences in State Trajectory Features

Figure 7 shows the OCR and NST values that we calculated. Note that all t-tests in this section and the following section were performed as HCs minus SZs. As such, a negative t-statistic means that SZs had higher values, and a positive t-statistic means that HCs had higher values. HCs switched states slightly more often than SZs (t = 3.74, p < 0.01). SZs spent significantly more time in states 0 (t = −3.65, p < 0.001) and 1 (t = −6.24, p < 0.001) than HCs. However, HCs spent significantly more time in states 2 (t = 6.24, p < 0.001) and 3 (t = 4.53, p < 0.001) than SZs.

**Figure 7.**
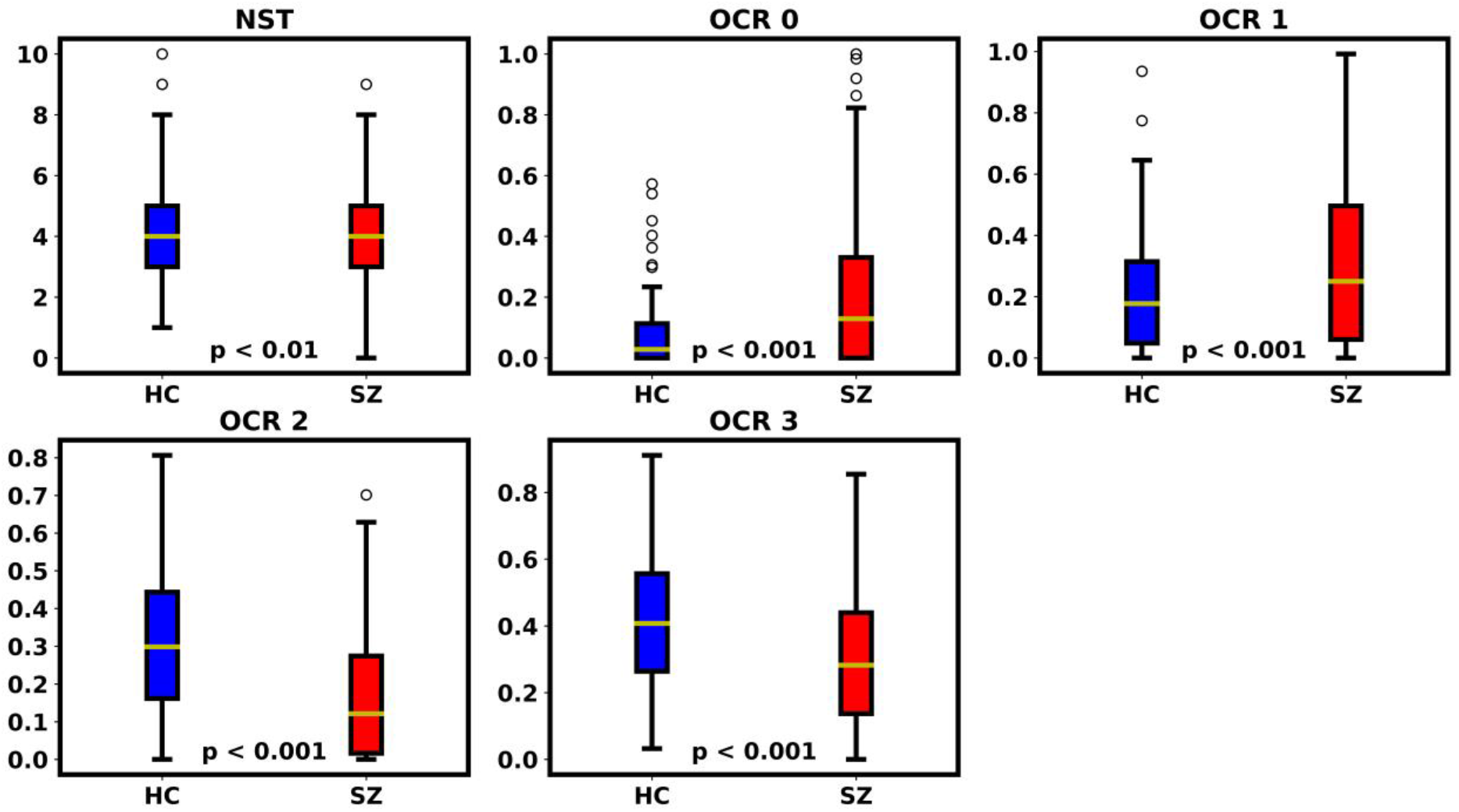
Number of State Transition and Occupancy Rate Features Per Class. The top left panel shows NST values for each class. The remaining panels show OCR values for the state numbers shown in their title. The leftmost (blue) and rightmost (red) boxplot in each panel shows feature values for HCs and SZs, respectively. FDR-corrected p-values resulting from the t-tests comparing HC and SZ values are shown at the bottom of each panel.

### Quantifying Class-Level Differences in Relevance Distributions

Figure 8 shows the EMD and KLD values for the relevance. The total EMD seems to be about the same for SZs and HCs. The total EMD indicates that HCs had slightly more (i.e., based on t-statistic but insignificant p-value) temporally concentrated relevance than SZs. However, based on the median EMD in Figure 8, it seems that the opposite is the case. This suggests that some SZs have more temporally concentrated relevance. Our KLD results are more consistent. HCs had significantly higher KLD range (t = 6.30, p < 0.001), standard deviation (t = 10.03, p < 0.001), median (t = 2.46, p < 0.05), mean (t = 3.23, p < 0.01), and max (t = 7.16, p < 0.001) values than SZs. This indicates that HCs had more change in the spatial distribution of importance across time points and that the magnitude of the change in spatial distribution varied more.

**Figure 8.**
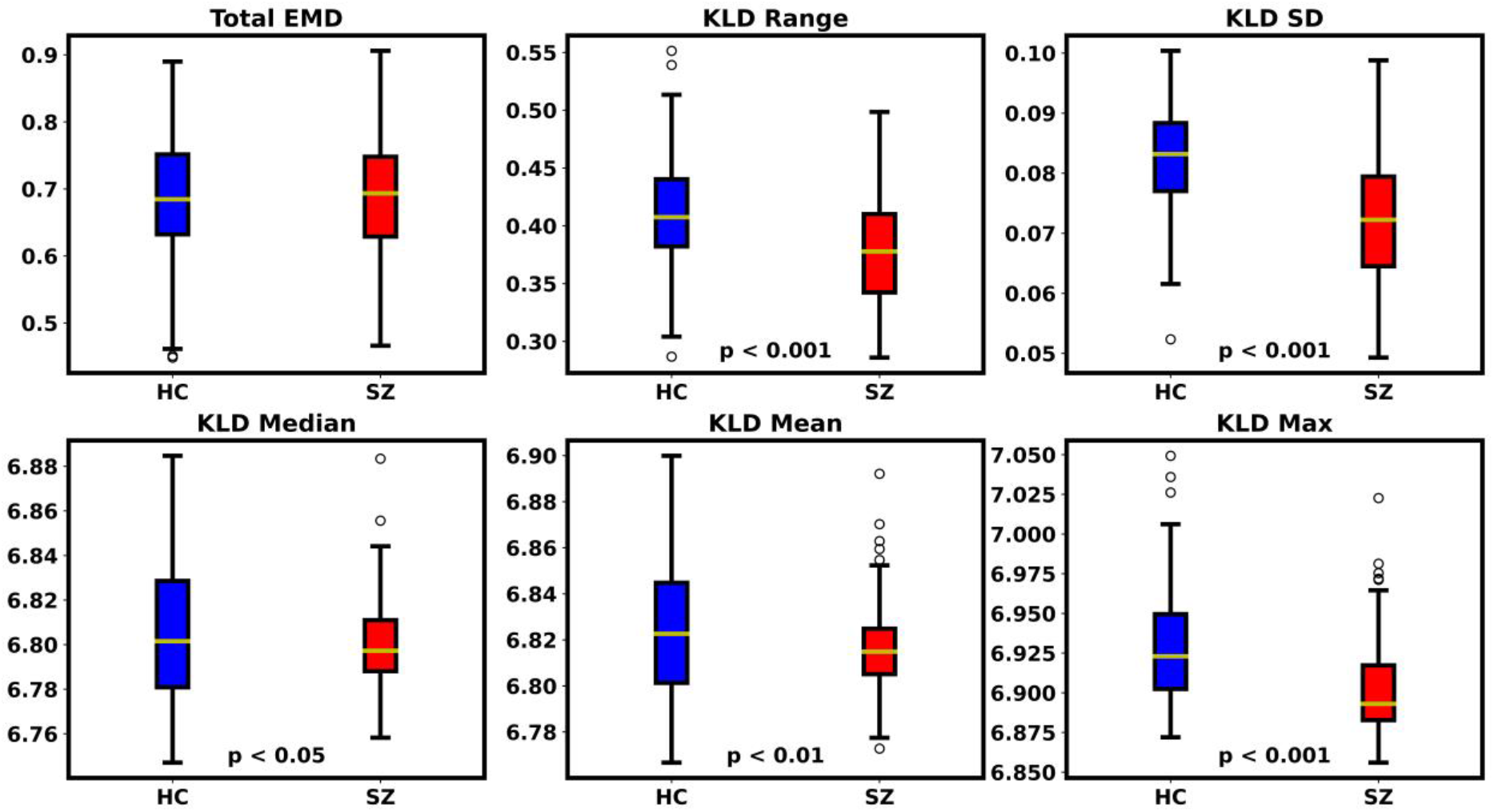
EMD and KLD Features Per Class. The top left panel shows total EMD values for each class. The remaining panels show KLD features shown in their title. The leftmost (blue) and rightmost (red) boxplot in each panel shows feature values for HCs and SZs, respectively. FDR-corrected p-values resulting from the t-tests comparing HC and SZ values are shown at the bottom of each panel. If no p-value is present, then the t-test had a non-significant result.

### Quantifying Relationships between State Trajectory Features and Relevance Distributions with Symptom Severity

Although we sought to identify relationships between the state-related features and relevance distribution features and symptom severity, there were no statistically significant relationships.

## DISCUSSION

In this study, we sought to (1) address a gap in existing machine learning-based dFNC analyses and (2) to provide new insights into the effects of SZ upon brain network interactions over time. Existing machine learning-based dFNC analyses have typically taken the form of classification, clustering, or clustering followed by classification. We developed a novel combination of an explainable classifier followed by a clustering algorithm to identify discriminatory patterns of dFNC and the durations for which those patterns are discriminatory. We then applied our approach within the context of schizophrenia, identifying novel patterns of SZ dFNC.

Our model test performance was well-above chance-level and higher than related studies on the same dataset (8,9), indicating that the patterns the model identified have the potential to be generalizable beyond our model and dataset. Additionally, we output explanations for the training data, for which the model had near perfect classification performance, and averaged explanations for participants across folds to reduce noise, obtain a more robust explanation, and increase the reliability of analysis results.

Importantly, SZs spent a median of 15% of their time with moderately negative VSN/SMN dFNC that differentiated them from HCs (i.e., state 0). Additionally, HCs spent about 20% of their time with strongly negative CBN/SMN and CBN/VSN activity that was highly important for differentiating them from SZs (i.e., state 1). SZs did spend some time in a state with slightly negative CBN/SMN (i.e., state 2). However, the median dFNC values for each state for SZs indicated that they typically did not have any negative CBN/VSN activity. As such, the interaction of the CBN and sensory networks seem to be particularly affected in SZ. These VSN, SMN, and CBN interactions are particularly interesting, as SZs tended to have less correlation between the CBN and sensory networks (i.e., VSN and SMN) but more negative correlation between sensory networks. The interactions of the sensory networks and the SCN also seemed to be affected by SZ. Namely, SZs spent around 10% of their time in a state with negative SCN/SMN and SCN/VSN dFNC (i.e., state 2). In contrast, HCs spent around 20% of their time in a state with even more strongly negative SCN/SMN and SCN/VSN dFNC (i.e., state 1). These combined findings suggest that SZs tend to have more frequent negative correlation between the VSN and SMN (83) but less interaction between the sensory networks and CBN (10,84) and SCN (45). Previous studies have also identified effects of SZ upon sensory network interactions (16) and the SCN (57).

Our approach also yielded useful insights into the temporal and spatial effects of SZ. Namely, based on our KLD relevance analysis, the features important for identifying HCs tended to vary more over time than SZs. This suggests that SZs could have reduced dynamism or variability in brain activity over time and fits well with existing discoveries (20). Based on our EMD relevance analysis, we also determined that the effects of SZ may be somewhat temporally concentrated. However, this finding is less certain. Some studies have found the effects of SZ upon ICs to be more temporally localized (46), while others have found the opposite to be the case in dFNC (9).

### Limitations and Future Work

Our analysis used a 40-second window size when extracting dFNC, which has been shown to be reasonable (65). However, the window size does affect the dFNC and our findings. We trained our model for multiple folds, output explanations for the training data, and averaged explanations across folds, to help increase the generalizability of our findings. However, future analyses might validate the reproducibility of our approach with different classifiers or SZ datasets. Additionally, we applied a hard clustering approach, and several dFNC clustering studies have highlighted the benefits of fuzzy clustering approaches (17,19,20). As such, in future iterations of our approach it might be helpful to investigate the use of fuzzy clustering methods.

## CONCLUSION

Many studies have analyzed rs-fMRI dFNC to characterize the effects of neurological and neuropsychiatric disorders upon brain activity. These studies have often applied classification or clustering approaches, and several studies have also performed clustering followed by classification. However, the use of explainable classifiers followed by clustering algorithms remains largely unexplored and presents opportunities for different insights into disorders. Namely, it enables improved insights into dynamics over classifiers and enhanced identification of discriminatory patterns over clustering approaches. In this study, we present such an approach, training an explainable deep learning model to differentiate between individuals with SZ and HCs from whole-brain dFNC data. We then cluster the resulting explanations to identify discriminatory states of brain activity. In particular, we uncover widespread effects of schizophrenia upon the interactions of the subcortical, sensory (i.e., visual and somatosensory), and cerebellar networks. We further extract several novel features to quantify different aspects of the classifier explanations, finding that individuals with SZ likely have reduced variability in overall brain activity and that the effects of SZ may also be temporally localized. Our proposed approach yields useful insights into the effects of SZ upon brain network dynamics. Moreover, it has the potential to provide novel insights into a variety of neurological and neuropsychiatric disorders in the future.

## REFERENCES

1. Sen B, Mueller B, Klimes-Dougan B, Cullen K, Parhi KK. Classification of Major Depressive Disorder from Resting-State fMRI. Proc Annu Int Conf IEEE Eng Med Biol Soc EMBS (2019)3511–3514. doi: 10.1109/EMBC.2019.8856453

2. Chun JY, Sendi MSE, Sui J, Zhi D, Calhoun VD. Visualizing Functional Network Connectivity Difference between Healthy Control and Major Depressive Disorder Using an Explainable Machine-learning Method. 2020 42nd Annual International Conference of the IEEE Engineering in Medicine & Biology Society (EMBC). (2020). p. 955–960 doi: 10.1109/BIBE50027.2020.00162

3. Rodriguez CI, Vergara VM, Davies S, Calhoun VD, Savage DD, Hamilton DA. Detection of prenatal alcohol exposure using machine learning classification of resting-state functional network connectivity data. Alcohol (2021) 93:25–34. doi: 10.1016/j.alcohol.2021.03.001

4. Challis E, Hurley P, Serra L, Bozzali M, Oliver S, Cercignani M. Gaussian process classification of Alzheimer’s disease and mild cognitive impairment from resting-state fMRI. Neuroimage (2015) doi: 10.1016/j.neuroimage.2015.02.037

5. Sendi MSE, Chun JY, Calhoun VD. Visualizing functional network connectivity difference between middle adult and older subjects using an explainable machine-learning method. Proceedings - IEEE 20th International Conference on Bioinformatics and Bioengineering, BIBE 2020. (2020). p. 955–960 doi: 10.1109/BIBE50027.2020.00162

6. Rashid B, Arbabshirani MR, Damaraju E, Cetin MS, Miller R, Pearlson GD, Calhoun VD. Classification of schizophrenia and bipolar patients using static and dynamic resting-state fMRI brain connectivity. Neuroimage (2016) 134:645–657. doi: 10.1016/j.neuroimage.2016.04.051

7. Deshpande G, Wang P, Rangaprakash D, Wilamowski B. Fully connected cascade artificial neural network architecture for attention deficit hyperactivity disorder classification from functional magnetic resonance imaging data. IEEE Trans Cybern (2015) 45:2668–2679. doi: 10.1109/TCYB.2014.2379621

8. Ellis CA, Miller RL, Calhoun VD. An Approach for Estimating Explanation Uncertainty in fMRI dFNC Classification. 2022 IEEE 22nd Int Conf Bioinforma Bioeng (2022)

9. Ellis CA, Miller RL, Calhoun VD. Towards Greater Neuroimaging Classification Transparency via the Integration of Explainability Methods and Confidence Estimation Approaches. Informatics Med Unlocked (2023) 37: doi: https://doi.org/10.1016/j.imu.2023.101176

10. Du Y, Fryer SL, Fu Z, Lin D, Sui J, Chen J, Damaraju E, Mennigen E, Stuart B, Mathalon DH, et al. Dynamic functional connectivity impairments in early schizophrenia and clinical high-risk for psychosis. Neuroimage (2018) 180:632–645. doi: 10.1016/j.neuroimage.2017.10.022.Dynamic

11. Sendi MSE, Zendehrouh E, Ellis CA, Liang Z, Fu Z, Mathalon DH, Ford JM, Preda A, van Erp TGM, Miller RL, et al. Aberrant Dynamic Functional Connectivity of Default Mode Network in Schizophrenia and Links to Symptom Severity. Front Neural Circuits (2021) 15:1–14. doi: 10.3389/fncir.2021.649417

12. Sendi MSE, Dini H, Bruni LE, Calhoun VD. Default mode network dynamic functional network connectivity predicts psychotic symptom severity. Proc Annu Int Conf IEEE Eng Med Biol Soc EMBS (2022) 2022-July:247–250. doi: 10.1109/EMBC48229.2022.9871542

13. Sendi MSE, Salat DH, Miller RL, Calhoun VD. Two-step clustering-based pipeline for big dynamic functional network connectivity data. Front Neurosci (2022) 16: doi: 10.3389/fnins.2022.895637

14. Rahaman A, Damaraju E, Turner JA, Erp TGM Van, Mathalon D, Muller B, Pearlson G, Calhoun VD. A novel method for tri-clustering dynamic functional network connectivity (dFNC) identifies significant schizophrenia effects across multiple states in distinct subgroups of individuals. bioRxiv (2020)

15. Li S, Hu N, Zhang W, Tao B, Dai J, Gong Y, Tan Y, Cai D, Lui S. Dysconnectivity of multiple brain networks in schizophrenia: A meta-analysis of resting-state functional connectivity. Front Psychiatry (2019) 10:1–11. doi: 10.3389/fpsyt.2019.00482

16. Salman MS, Du Y, Calhoun VD. Identifying FMRI dynamic connectivity states using affinity propagation clustering method: Application to schizophrenia. ICASSP, IEEE Int Conf Acoust Speech Signal Process - Proc (2017)904–908. doi: 10.1109/ICASSP.2017.7952287

17. Ellis CA, Miller RL, Calhoun VD. A Novel Explainable Fuzzy Clustering Approach for fMRI Dynamic Functional Network Connectivity Analysis. bioRxiv. (2023)

18. Ellis CA, Miller RL, Calhoun VD. Explainable Fuzzy Clustering Framework Reveals Divergent Default Mode Network Connectivity Dynamics in Schizophrenia. bioRxiv (2023)

19. Abrol A, Damaraju E, Miller RL, Stephen J, Claus E, Mayer A, Calhoun VD. Replicability of time-varying connectivity patterns in large resting state fMRI samples. Neuroimage (2017) 163:160–176. doi: 10.1016/j.neuroimage.2017.09.020.Replicability

20. Miller RL, Yaesoubi M, Turner JA, Mathalon D, Preda A, Pearlson G, Adali T, Calhoun VD. Higher dimensional meta-state analysis reveals reduced resting fMRI connectivity dynamism in schizophrenia patients. PLoS One (2016) 11:1–24. doi: 10.1371/journal.pone.0149849

21. Du Y, Pearlson GD, Yu Q, He H, Lin D, Sui J, Wu L, Calhoun VD. Interaction among subsystems within default mode network diminished in schizophrenia patients: A dynamic connectivity approach. Schizophr Res (2016) 170:55–65. doi: 10.1016/j.schres.2015.11.021

22. Ellis CA, Sendi MSE, Miller RL, Calhoun VD. An Unsupervised Feature Learning Approach for Elucidating Hidden Dynamics in rs-fMRI Functional Network Connectivity. 2022 44th Annual International Conference of the IEEE Engineering in Medicine & Biology Society (EMBC). IEEE (2022). p. 4449–4452

23. Sendi MSE, Pearlson GD, Mathalon DH, Ford JM, Preda A, Erp TGM Van, Calhoun VD. Multiple overlapping dynamic patterns of the visual sensory network in schizophrenia. Schizophr Res (2021) 228:103–111. doi: 10.1016/j.schres.2020.11.055

24. Damaraju E, Allen EA, Belger A, Ford JM, McEwen S, Mathalon DH, Mueller BA, Pearlson GD, Potkin SG, Preda A, et al. Dynamic functional connectivity analysis reveals transient states of dysconnectivity in schizophrenia. NeuroImage Clin (2014) 5:298–308. doi: 10.1016/j.nicl.2014.07.003

25. Fu Z, Iraji A, Turner JA, Sui J, Miller R, Pearlson GD, Calhoun VD. Dynamic state with covarying brain activity-connectivity: On the pathophysiology of schizophrenia. Neuroimage (2021) 224:117385. doi: 10.1016/j.neuroimage.2020.117385

26. Sendi MSE, Zendehrouh E, Fu Z, Liu J, Du Y, Mormino E, Salat DH, Calhoun VD, Miller RL. Disrupted Dynamic Functional Network Connectivity Among Cognitive Control Networks in the Progression of Alzheimer’s Disease. Brain Connect (2021)1–25. doi: 10.1089/brain.2020.0847

27. Sendi MSE, Zendehrouh E, Ellis CA, Chen J, Miller RL, Mormino EC, Salat DH, Calhoun VD. The link between brain functional network connectivity and genetic risk of Alzheimer’s disease. bioRxiv (2021) doi: 10.1002/alz.050101

28. Sendi MSE, Ellis CA, Milller RL, Salat DH, Calhoun VD. The relationship between dynamic functional network connectivity and spatial orientation in healthy young adults. bioRxiv (2021)

29. Sendi MSE, Zendehrouh E, Fu Z, Mahmoudi B, Miller RL, Calhoun VD. A Machine Learning Model for Exploring Aberrant Functional Network Connectivity Transition in Schizophrenia. Proceedings of the IEEE Southwest Symposium on Image Analysis and Interpretation. (2020). p. 112–115 doi: 10.1109/SSIAI49293.2020.9094620

30. Sendi MSE, Member S, Kanta V, Inman CS, Manns JR, Hamann S, Gross RE, Willie JT, Mahmoudi B. Amygdala Stimulation Leads to Functional Network Connectivity State Transitions in the Hippocampus. 42nd Annual International Conference of the IEEE Engineering in Medicine and Biology Society (EMBC). (2020). p. 3625–3628

31. Gawne TJ, Overbeek GJ, Killen JF, Reid MA, Kraguljac N V., Denney TS, Ellis CA, Lahti AC. A multimodal magnetoencephalography 7 T fMRI and 7 T proton MR spectroscopy study in first episode psychosis. npj Schizophr (2020) 6:1–9. doi: 10.1038/s41537-020-00113-4

32. Ellis CA, Sattiraju A, Miller R, Calhoun V. Examining Effects of Schizophrenia on EEG with Explainable Deep Learning Models. 2022 IEEE 22nd International Conference on Bioinformatics and Bioengineering (BIBE). (2022)

33. Ellis CA, Sattiraju A, Miller R, Calhoun V. Examining Reproducibility of EEG Schizophrenia Biomarkers Across Explainable Machine Learning Models. bioRxiv. (2022)

34. Espinoza FA, Vergara VM, Damaraju E, Henke KG, Faghiri A, Turner JA, Belger AA, Ford JM, McEwen SC, Mathalon DH, et al. Characterizing whole brain temporal variation of functional connectivity via zero and first order derivatives of sliding window correlations. Front Neurosci (2019) 13:1–13. doi: 10.3389/fnins.2019.00634

35. Calhoun VD. Dynamic connectivity states estimated from resting fMRI Identify differences among Schizophrenia, bipolar disorder, and healthy control subjects. (2014) 8:1–13. doi: 10.3389/fnhum.2014.00897

36. Zhang L. EEG Signals Classification Using Machine Learning for the Identification and Diagnosis of Schizophrenia. Proc Annu Int Conf IEEE Eng Med Biol Soc EMBS (2019)4521–4524. doi: 10.1109/EMBC.2019.8857946

37. Shim M, Hwang HJ, Kim DW, Lee SH, Im CH. Machine-learning-based diagnosis of schizophrenia using combined sensor-level and source-level EEG features. Schizophr Res (2016) 176:314–319. doi: 10.1016/j.schres.2016.05.007

38. Bliksted V, Frith C, Videbech P, Fagerlund B, Emborg C, Simonsen A, Roepstorff A, Campbell-Meiklejohn D. Hyper-and Hypomentalizing in Patients with First-Episode Schizophrenia: FMRI and Behavioral Studies. Schizophr Bull (2019) 45:377–385. doi: 10.1093/schbul/sby027

39. Ebisch SJH, Gallese V, Salone A, Martinotti G, di Iorio G, Mantini D, Perrucci MG, Romani GL, Di Giannantonio M, Northoff G. Disrupted relationship between “resting state” connectivity and task-evoked activity during social perception in schizophrenia. Schizophr Res (2018) 193:370–376. doi: 10.1016/j.schres.2017.07.020

40. Shukla DK, Wijtenburg SA, Chen H, Chiappelli JJ, Kochunov P, Hong LE, Rowland LM. Anterior cingulate glutamate and GABA associations on functional connectivity in schizophrenia. Schizophr Bull (2019) 45:647–658. doi: 10.1093/schbul/sby075

41. Hare SM, Ford JM, Mathalon DH, Damaraju E, Bustillo J, Belger A, Lee HJ, Mueller BA, Lim KO, Brown GG, et al. Salience-default mode functional network connectivity linked to positive and negative symptoms of schizophrenia. Schizophr Bull (2019) 45:892–901. doi: 10.1093/schbul/sby112

42. Kottaram A, Johnston LA, Cocchi L, Ganella EP, Everall I, Pantelis C, Kotagiri R, Zalesky A. Brain network dynamics in schizophrenia: Reduced dynamism of the default mode network. Hum Brain Mapp (2019) 40:2212–2228. doi: 10.1002/hbm.24519

43. Chand GB, Thakuri DS, Soni B, Kingshighway Blvd St Louis S. Disrupted controlling mechanism of salience network on default-mode network and central-executive network in schizophrenia. bioRxiv (2021)1–19. https://doi.org/10.1101/2021.12.03.471183

44. Whitfield-Gabrieli S, Thermenos HW, Milanovic S, Tsuang MT, Faraone S V., McCarley RW, Shenton ME, Green AI, Nieto-Castanon A, LaViolette P, et al. Hyperactivity and hyperconnectivity of the default network in schizophrenia and in first-degree relatives of persons with schizophrenia. Proc Natl Acad Sci U S A (2009) 106:1279–1284. doi: 10.1073/pnas.0809141106

45. Yu Q, Calhoun VD. Resting-State Functional Network Disturbances in Schizophrenia. Brain Netw Dysfunct Neuropsychiatr Illn (2021)187–215. doi: 10.1007/978-3-030-59797-9_10

46. Rahman M, Lewis N, Fedorov A, Calhoun V, Rahman M, Mahmood U, Lewis N, Gazula H. Interpreting models interpreting brain dynamics. Sci Rep (2022) 12:1–16.

47. Sanfratello L, Houck J, Calhoun VD. Dynamic Functional Network Connectivity In Schizophrenia With MEG And fMRI, Do Different Time Scales Tell A Different Story? Brain Connect (2019) doi: 10.1089/brain.2018.0608

48. Zendehrouh E, Sendi MSE, Sui J, Fu Z, Zhi D, Lv L, Ma X, Ke Q, Li X, Wang C, et al. Aberrant Functional Network Connectivity Transition Probability in Major Depressive Disorder. 42nd Annual International Conference of the IEEE Engineering in Medicine and Biology Society (EMBC). Montreal, QC, Canada: IEEE (2020). p. 1493–1496

49. Wu X jie, Zeng LL, Shen H, Yuan L, Qin J, Zhang P, Hu D. Functional network connectivity alterations in schizophrenia and depression. Psychiatry Res - Neuroimaging (2017) 263:113–120. doi: 10.1016/j.pscychresns.2017.03.012

50. Dini H, Sendi MSE, Sui J, Fu Z, Espinoza R, Narr KL, Qi S, Abbott CC, van Rooij SJH, Riva-Posse P, et al. Dynamic Functional Connectivity Predicts Treatment Response to Electroconvulsive Therapy in Major Depressive Disorder. Front Hum Neurosci (2021) 15:1–11. doi: 10.3389/fnhum.2021.689488

51. Sendi MSE, Zendehrouh E, Miller RL, Fu Z, Du Y, Liu J, Mormino EC, Salat DH, Calhoun VD. Alzheimer’s Disease Projection From Normal to Mild Dementia Reflected in Functional Network Connectivity: A Longitudinal Study. Front Neural Circuits (2021) 14: doi: 10.3389/fncir.2020.593263

52. Fu Z, Sui J, Turner JA, Du Y, Assaf M, Pearlson GD, Calhoun VD. Dynamic functional network reconfiguration underlying the pathophysiology of schizophrenia and autism spectrum disorder. Hum Brain Mapp (2021) 42:80–94. doi: 10.1002/hbm.25205

53. Ellis CA, Sancho ML, Miller R, Calhoun V. Exploring Relationships between Functional Network Connectivity and Cognition with an Explainable Clustering Approach. 2022 IEEE 22nd International Conference on Bioinformatics and Bioengineering (BIBE). IEEE (2022). p. 23–26 doi: 10.1109/BIBE55377.2022.00066

54. Ellis CA, Sendi MSE, Geenjaar EPT, Plis SM, Miller RL, Calhoun VD. Algorithm-Agnostic Explainability for Unsupervised Clustering. (2021)1–22. http://arxiv.org/abs/2105.08053

55. Damaraju E, Allen EA, Belger A, Ford JM, McEwen S, Mathalon DH, Mueller BA, Pearlson GD, Potkin SG, Preda A, et al. Dynamic functional connectivity analysis reveals transient states of dysconnectivity in schizophrenia. NeuroImage Clin (2014) 5:298–308. doi: 10.1016/j.nicl.2014.07.003

56. Fu Z, Caprihan A, Chen J, Du Y, Adair JC, Sui J, Rosenberg GA, Calhoun VD. Altered static and dynamic functional network connectivity in Alzheimer’s disease and subcortical ischemic vascular disease: shared and specific brain connectivity abnormalities. Hum Brain Mapp (2019) 40:3203–3221. doi: 10.1002/hbm.24591

57. Sun Y, Collinson SL, Suckling J, Sim K. Dynamic reorganization of functional connectivity reveals abnormal temporal efficiency in schizophrenia. Schizophr Bull (2019) 45:659–669. doi: 10.1093/schbul/sby077

58. Sendi MSE, Zendehrouh E, Fu Z, Liu J, Du Y, Mormino E, Salat DH, Calhoun VD, Miller RL. Disrupted Dynamic Functional Network Connectivity Among Cognitive Control Networks in the Progression of Alzheimer’s Disease. Brain Connect (2021)1–25. doi: 10.1089/brain.2020.0847

59. Ellis CA, Sendi MSE, Zhang R, Carbajal DA, Wang MD, Miller L, Calhoun VD. Novel Methods for Elucidating Modality Importance in Multimodal Electrophysiology Classifiers. bioRxiv (2022)

60. Ellis CA, Miller RL, Calhoun VD. Neuropsychiatric Disorder Subtyping Via Clustered Deep Learning Classifier Explanations. bioRxiv. (2022). p. 12–15

61. Amunts K, Mohlberg H, Bludau S, Zilles K. Julich-Brain: A 3D probabilistic atlas of the human brain’s cytoarchitecture. Science (80-) (2020) 369:988–992. doi: 10.1126/science.abb4588

62. van Erp TGM, Preda A, Turner JA, Callahan S, Calhoun VD, Bustillo JR, Lim KO, Mueller B, Brown GG, Vaidya JG, et al. Neuropsychological profile in adult schizophrenia measured with the CMINDS. Psychiatry Res (2015) 230:826–834. doi: 10.1016/j.psychres.2015.10.028.Neuropsychological

63. Ellis CA., Miller RL., Calhoun VD. Identifying Neuropsychiatric Disorder Subtypes and Subtype-Dependent Variation in Diagnostic Deep Learning Classifier Performance. bioRxiv (2022)2–5.

64. Kay SR, Fiszbein A, Opler LA. The positive and negative syndrome scale (PANSS) for schizophrenia. Schizophr Bull (1987) 13:261–276. doi: 10.1093/schbul/13.2.261

65. Fu Z, Du Y, Calhoun VD. The Dynamic Functional Network Connectivity Analysis Framework. Engineering (2019) 5:190–193. doi: 10.1016/j.eng.2018.10.001

66. Chollet F. Keras. GitHub (2015) https://github.com/fchollet/keras

67. Kingma DP, Ba J. Adam: A method for stochastic optimization. Proceedings of the 3rd International Conference on Learning Representations (ICLR). (2015)

68. Glorot X, Bengio Y. Understanding the difficulty of training deep feedforward neural networks. J Mach Learn Res (2010) 9:249–256.

69. He K, Zhang X, Ren S, Sun J. Delving deep into rectifiers: Surpassing human-level performance on imagenet classification. Proc IEEE Int Conf Comput Vis (2015) 2015 Inter:1026–1034. doi: 10.1109/ICCV.2015.123

70. Samek W, Montavon G, Vedaldi A, Hansen LK, Müller K-R eds. Explainable AI: Interpreting, Explaining and Visualizing Deep Learning. Cham: Springer International Publishing (2019). doi: 10.1007/978-3-030-28954-6

71. Bach S, Binder A, Montavon G, Klauschen F, Müller KR, Samek W. On pixel-wise explanations for non-linear classifier decisions by layer-wise relevance propagation. PLoS One (2015) 10: doi: 10.1371/journal.pone.0130140

72. Samek W, Binder A, Montavon G, Lapuschkin S, Müller KR. Evaluating the visualization of what a deep neural network has learned. IEEE Trans Neural Networks Learn Syst (2017) 28:2660–2673. doi: 10.1109/TNNLS.2016.2599820

73. Yan W, Plis S, Calhoun VD, Liu S, Jiang R, Jiang T-Z, Sui J. Discriminating Schizophrenia From Normal Controls Using Resting State Functional Network Connectivity: A Deep Neural Network and Layer-wise Relevance Propagation Method. IEEE INTERNATIONAL WORKSHOP ON MACHINE LEARNING FOR SIGNAL PROCESSING. (2017)

74. Sturm I, Lapuschkin S, Samek W, Müller KR. Interpretable deep neural networks for single-trial EEG classification. J Neurosci Methods (2016) 274:141–145. doi: 10.1016/j.jneumeth.2016.10.008

75. Ellis CA, Miller RL, Calhoun VD A Systematic Approach for Explaining Time and Frequency Features Extracted by Convolutional Neural Networks From Raw Electroencephalography Data. Front Neuroinform (2022) 16:1–11. doi: 10.3389/fninf.2022.872035

76. Ellis CA, Sendi MS, Willie JT, Mahmoudi B. Hierarchical Neural Network with Layer-wise Relevance Propagation for Interpretable Multiclass Neural State Classification. 10th International IEEE/EMBS Conference on Neural Engineering (NER). (2021). p. 18–21

77. Thomas AW, Heekeren HR, Müller K-R, Samek W. Analyzing Neuroimaging Data Through Recurrent Deep Learning Models. (2018) http://arxiv.org/abs/1810.09945

78. Ellis CA, Miller RL, Calhoun VD A Model Visualization-based Approach for Insight into Waveforms and Spectra Learned by CNNs. Proceedings of the Annual International Conference of the IEEE Engineering in Medicine and Biology Society, EMBS. IEEE (2022). p. 1643–1646 doi: 10.1109/EMBC48229.2022.9871414

79. Ellis CA, Carbajal DA, Zhang R, Miller RL, Calhoun VD, Wang MD. An Explainable Deep Learning Approach for Multimodal Electrophysiology Classification. bioRxiv (2021)12–15.

80. Mayor-Torres JM, Medina-DeVilliers S, Clarkson T, Lerner MD, Riccardi G. Evaluation of Interpretability for Deep Learning algorithms in EEG Emotion Recognition: A case study in Autism. (2021)1–12. http://arxiv.org/abs/2111.13208

81. Rousseeuw PJ. Silhouettes: A graphical aid to the interpretation and validation of cluster analysis. J Comput Appl Math (1987) 20:53–65. doi: 10.1016/0377-0427(87)90125-7

82. Benjamini Y, Hochberg Y. Controlling the False Discovery Rate: A Practical and Powerful Approach to Multiple Testing. J R Stat Soc Ser B (1995) 57:289–300. doi: 10.1111/j.2517-6161.1995.tb02031.x

83. Chen X, Duan M, Xie Q, Lai Y, Dong L, Cao W, Yao D, Luo C. Functional disconnection between the visual cortex and the sensorimotor cortex suggests a potential mechanism for self-disorder in schizophrenia. Schizophr Res (2014) 166:151–157. doi: 10.1016/j.schres.2015.06.014

84. Liang M, Zhou Y, Jiang T, Liu Z, Tian L, Liu H, Hao Y. Widespread functional disconnectivity in schizophrenia with resting-state functional magnetic resonance imaging. Neuroreport (2006) 17:209–213. doi: 10.1097/01.wnr.0000198434.06518.b8

